## Supplementary files for "Sending mixed signals: convergent iridescence and divergent chemical signals in sympatric sister-species of Amazonian butterflies"

**Supporting Information Text 1: Morpho wing reflectance measurement protocols**

The reflectance of the right anterior wings of 80 *Morpho* individuals was measured at different angles of illumination using a spectrometer (AvaSpec-ULS2048CL-EVO-RS, Avantes) coupled with a deuterium halogen light source (AvaLight-DH-S-BAL, Avantes) and two optical fibers (FCR-7UVIR200-2-1.5X100 and FC-UVIR200-2-1.5X100, Avantes) supported by an AFH-15 Angled Fibre Holder (Avantes) (Fig. S1.).

First, we measured the specular reflectance. Specularity refers to the reflectance in the direction symmetrical to the illumination relative to the normal of a surface (*i.e.* the vector perpendicular to the surface of an object): it is the expected direction of maximal reflectance of a perfect mirror. (Fig. S2.). We used a total of 13 specular combinations to quantify the hue variation within samples due to iridescence. First, we positioned the two fibers vertically, *i.e* their common axis was perpendicular to the surface of the wings (forming a 0° angle) to measure the wing reflectance at the normal of the wing. Then, we jointly tilted the two fibers symmetrically to the normal in the proximo-distal plane and measured the reflectance of the wings at (i) a 15° angle from the normal, (ii) a 30° angle from the normal and (iii) a 45° angle from the normal. The illumination and observation fibers were then switched and the 3 specular measures were repeated under a new illumination side, allowing the quantification of the specular reflectance of the proximo-distal plane at 6 different angles of illumination in total. The same protocol was applied by positioning the two fibers symmetrically to the normal in the antero-posterior plane and measuring the reflectance of the wings at 6 different angles of illumination as well.

We then quantified the variation of brightness of the wings using a "tilt set-up" (Fig. S3)(1).

We symmetrically positioned the two fibers 30° to the normal of the wings, forming a constant angular span of 60° between the two fibers in the proximo-distal plane of the wings. While keeping this fixed position between the two fibers, we measured the wing reflectance when tilting the fibers (i) 15° toward the normal of the wings and (ii) 15° away from the normal of the wings. The illumination and observation fibers were then switched and 2 additional measurements were performed on the proximo-distal plane of the wings. The same protocol was applied by positioning the two fibers in the antero-posterior plane and measuring the reflectance of the wings at 4 new different angles of illumination. This protocol allowed for the quantification of 8 new reflectance spectra quantifying iridescence for each sample.

Note that for both methods of measurement, we measured a black and a white reference every time the angle of illumination changed.

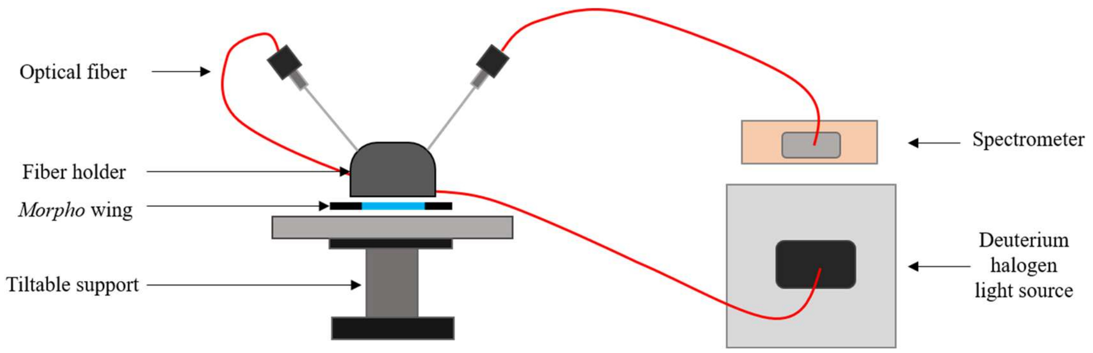


**S1 Fig** Illustration of the set-up used to measure the reflectance of the wings of *Morpho* butterflies at different angles of illumination and observation. We used a fiber holder to precisely control the angle between the two fibers. We ensured the measured surface was flat with a support holding the wings.

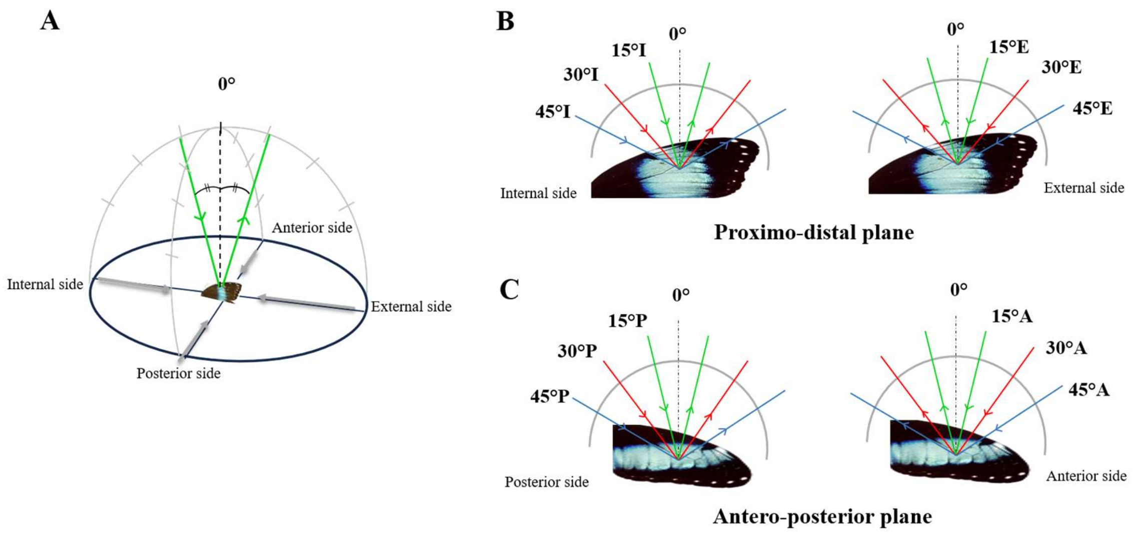


**S2 Fig** Scheme of the specular angles of illumination used to measure the reflectance of the wings. (A) shows the coverage of the 13 angles of illumination measured for each wing. The angles analyzed on the proximo-distal plane are represented in (B) and the angles analyzed on the antero-posterior plane are shown in (C). Each letter used to describe the angles (I, E, P, A) refer to the side the light was directed to (Internal, External, Posterior and Anterior side of wings, respectively).

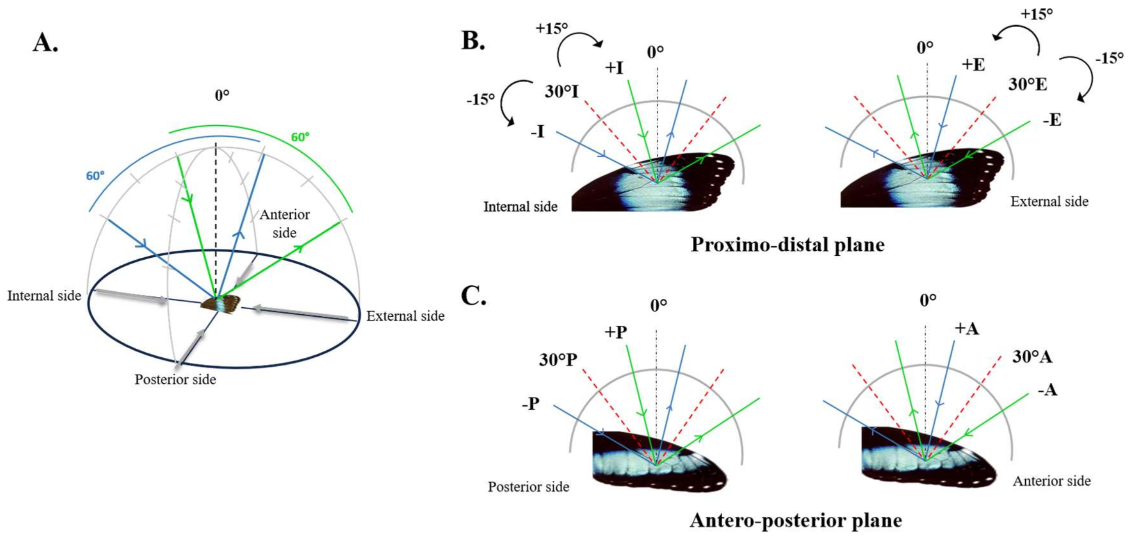


**S3 Fig** Scheme of the angles measured in the tilt set-up. The red dotted lines represent the specular 30° angle measured in the "specular" set-up. The angular span between the two fibers is kept the same and the two fibers are tilted toward the normal of the wings (annotated with +) or away from it (annotated with -), allowing the additional measurement of the wing's reflectance. This operation is repeated on the Internal, External, Anterior and Posterior sides (I, E, A and P respectively) of the wings.

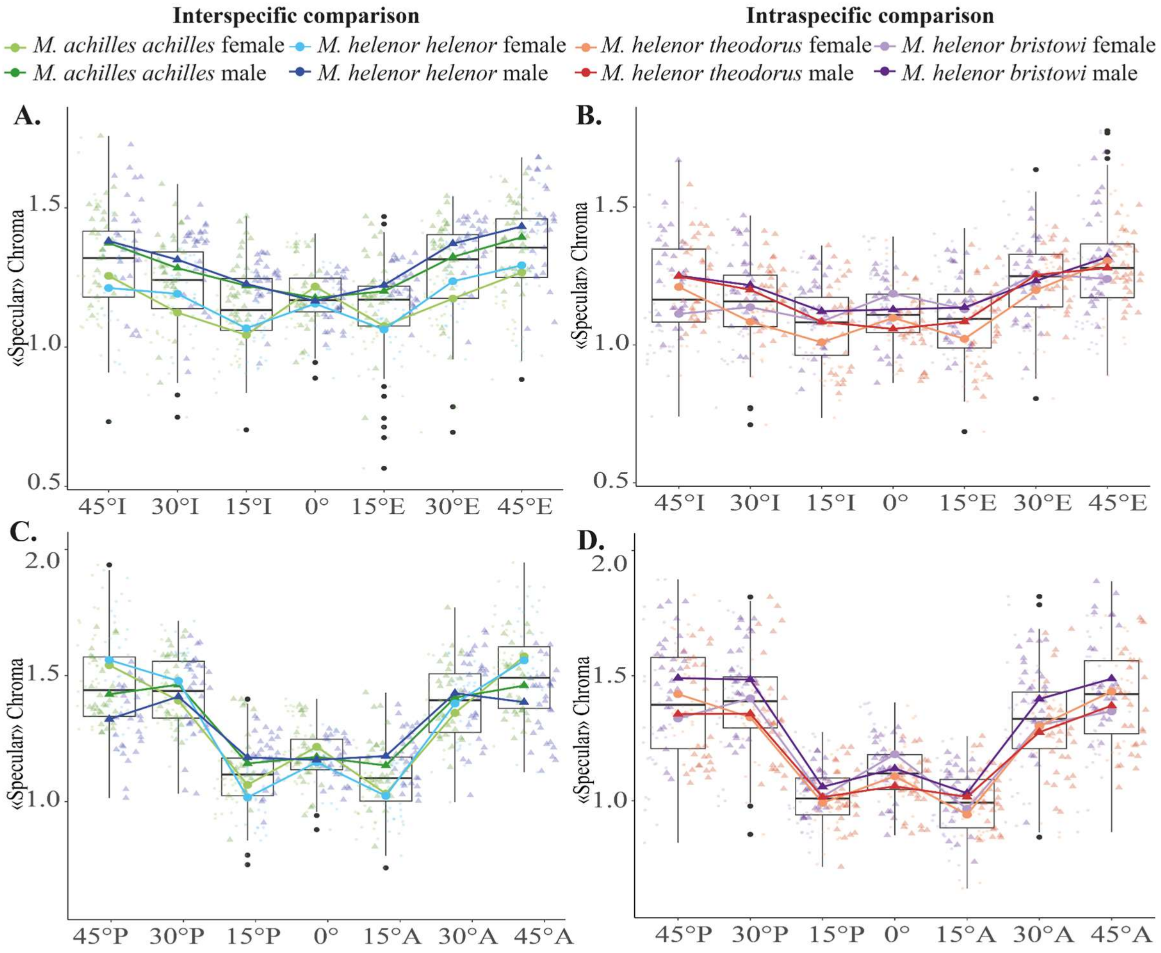


E.

| | Chroma specular EI French Guiana | Chroma specular EI Ecuador | Chroma specular AP French Guiana | Chroma specular AP Ecuador |
|---|---|---|---|---|
| **Sex** | **0.001***** | **0.001***** | 0.066 | **0.007**** |
| **Species** | 0.143 | **0.005**** | 0.241 | **0.001***** |
| **Angle** | **0.001***** | **0.001***** | **0.001***** | **0.001***** |
| **Sex:Species** | 0.682 | 0.836 | 0.379 | **0.001***** |
| **Species:Angle** | 0.137 | **0.003**** | 0.348 | 0.164 |

**S4 Fig** Variation of the chroma parameter measured on the proximo-distal plane (A and B) and anteroposterior plane (C and D) calculated from the wing reflectance measured with "Specular" set-up. The chroma parameter was calculated for the sympatric *M. h. helenor* and *M. a. achilles* (first column in green and blue) and for the allopatric *M. h. theodorus* and *M. h. bristowi* (second column in orange and purple). The results of the permutation-based ANOVAs performed in order to test whether the sex, the taxa or the angle of illumination have an effect on the estimated chroma are shown in (E). The chroma parameter is a proxy describing the intensity of the color reflected by the dorsal side of *Morpho* wings. As for every optical parameter measured, chroma is significantly different at every angle of illumination suggesting that variations of chroma can be observed on the wings of *Morpho* butterflies (iridescence). Overall, we can see on every graph that the colors measured at a wide specular angle (30° to 45° in every tested direction) have the highest chroma, suggesting the existence of a more intense color signal at extreme angles of illumination. On the proximo-distal plane (A and B), chroma is significantly different between males and females. The graphs show that males tend to be more saturated than females, especially at extreme angles. On the anteroposterior plane, especially among the sympatric *Morpho* species analysis (C), the chroma of males is overall higher than the female's, except at 45° angles of illumination where it drops for males but increases for females. The chroma parameter is always significantly different between *M. h. bristowi* and *M. h. theodorus* subspecies on both planes of illumination (B and D), and is also similar between sympatric *M. h. helenor* and *M. a. achilles* (A and C), consistent with convergence of the intensity of color saturation between sympatric species.

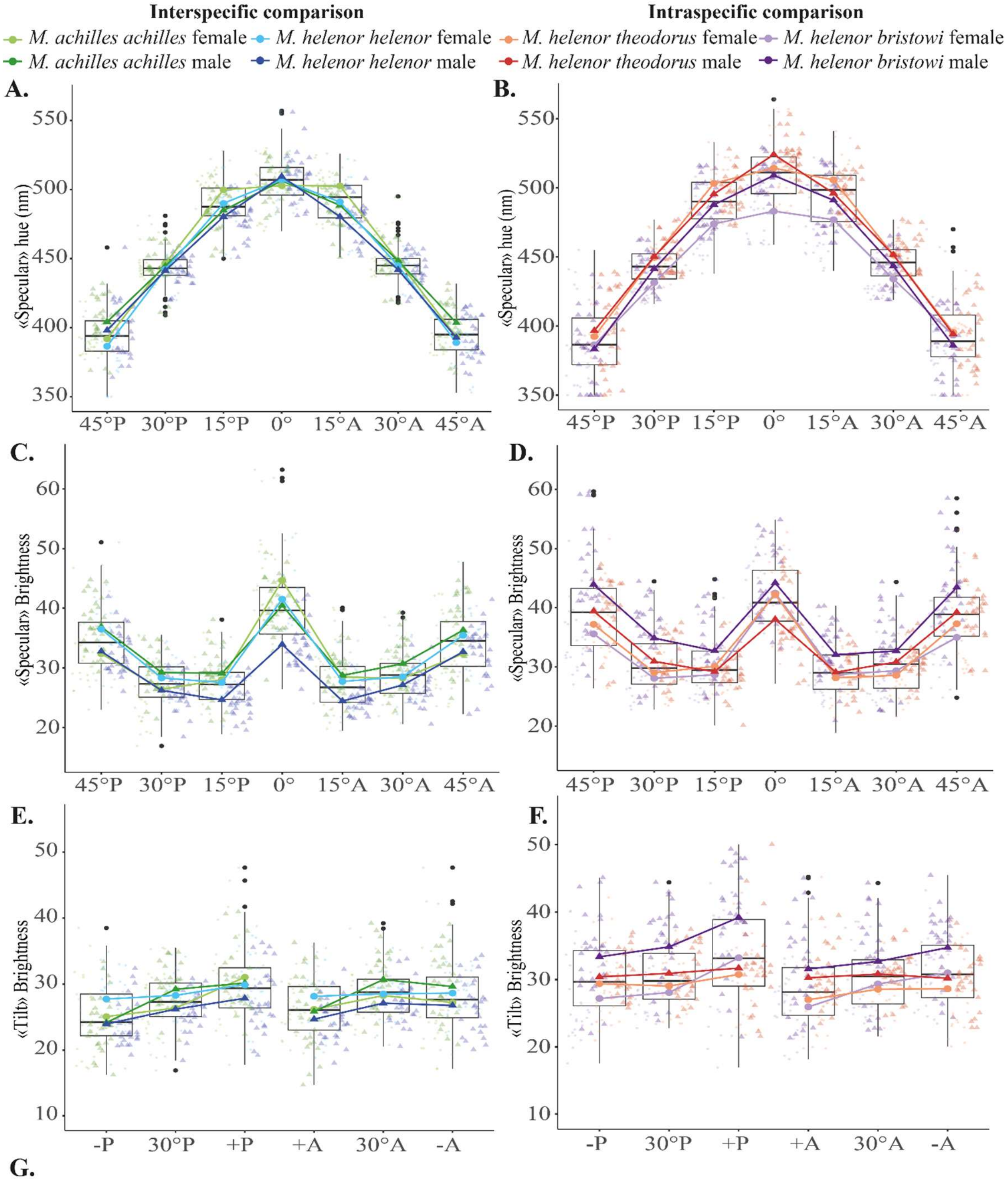


G.

| | Hue Specular French Guiana | Hue Specular Ecuador | Brightness Specular French Guiana | Brightness Specular Ecuador | Brightness « 30° tilt » French Guiana | Brightness « 30° tilt » Ecuador |
|---|---|---|---|---|---|---|
| **Sex** | 0.417 | **0.006**** | **0.012*** | **0.001***** | **0.031*** | **0.001***** |
| **Species** | **0.001***** | **0.001***** | **0.001***** | **0.001***** | 0.243 | **0.002**** |
| **Angle** | **0.001***** | **0.001***** | **0.001***** | **0.001***** | **0.001***** | **0.001***** |
| **Sex:Species** | 0.803 | **0.004***** | **0.001***** | **0.001***** | **0.001***** | **0.001***** |
| **Species:Angle** | **0.043*** | 0.099 | **0.001***** | 0.627 | 0.099 | **0.005***** |

**S5 Fig** Variations in hue (A and B) and brightness (C and D) calculated from the wing reflectance measured with the "specular" set-up, and variations of brightness calculated from the wing reflectance measured from the "tilt" set-up (E and F). Differences in hue and brightness of the anteroposterior plane of the wings are shown for both the interspecific (first column in green and blue), and intraspecific comparisons (second column in orange and purple). The results of the permutation-based ANOVAs performed in order to test whether the sex, the taxa or the angle of illumination, have an effect on the estimated hue and brightness are shown in (G). Similarly to the proximo-distal data presented in the main text (Figure 2), the analysis of the reflectance of *Morpho* wings on their anteroposterior plane shows that this plane is also iridescent as significant variations of hue and brightness are measured at different angles. The effect of sex on the variation of brightness is always significant no matter the pair tested or the method of measurement: the brightness of *Morpho* wings on this plane is thus sexually dimorphic (C, D, E and F) as was found on the proximo-distal plane. However, the "tilt" wing reflectance measurements of the brightness (E and F) are not as straightforward as the measurements taken on the proximo-distal plane showing that males were brighter than females (Fig. 2.F and 2.G). Here we observe that allopatric males are indeed brighter than females (F), but the Amazonian males from French Guiana are not as clearly different from their respective females (E). The difference of brightness between males and females in those two localities are also less important than the differences of brightness found between males and females measured on the proximo-distal plane (Fig. 2.F and 2.G). This difference of brightness could be explained by the physical structures of the scales that could potentially better reflect light intensity on the proximo-distal plane than on the anteroposterior plane, generating more important shifts in brightness during a flapping flight motion. Finally, a significant effect of hue was found between the two allopatric populations of *M. helenor* (B) and between the two sympatric *Morpho* species (A). Conversely, no hue variation was found between the sympatric *Morpho* species on the proximo-distal plane (Figure 2.B). Nevertheless, divergence in hue was found to be more important between the allopatric *M. helenor* subspecies than between the sympatric *M. h.*

*helenor* and *M. a. achilles*, consistent with stronger convergence in coloration between sympatric species.

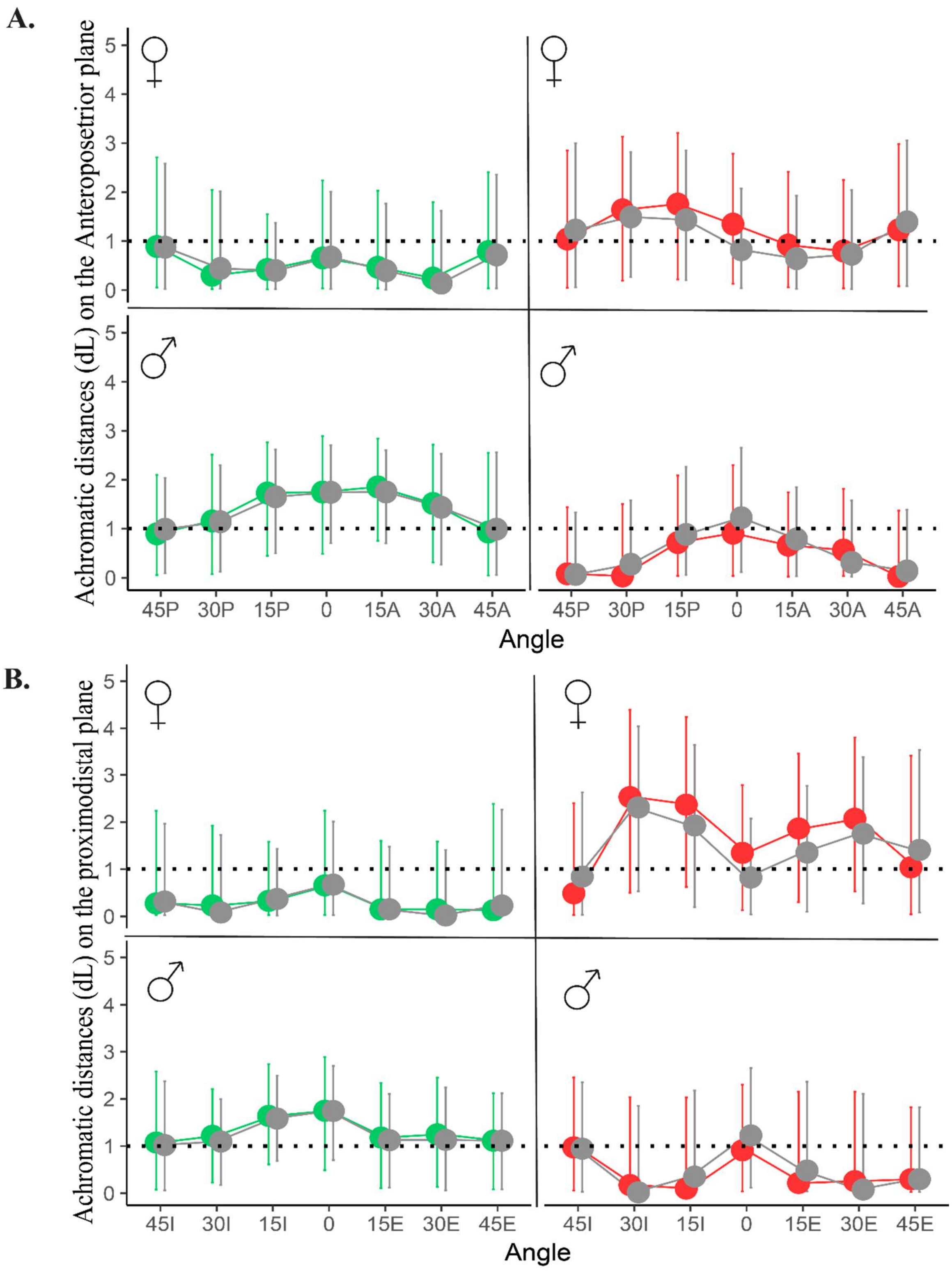


**S6 Fig** Achromatic distances (i.e. the visual discrimination rate of brightness by a visual model) of the wing reflectance measured with the "Specular" set-up on (A) the anteroposterior plane and

(B) the proximodistal plane. We used visual modeling to calculate the achromatic contrast found between the blue coloration of the two allopatric population of *M.helenor* sampled in Ecuador, as seen by a *Morpho* visual system (in red) and between the blue coloration of two species of Morphos from French Guiana, as seen by a *Morpho* visual system (in green). We also added the achromatic contrast of the *Morpho* wings as seen by the visual system of an avian predator (in grey). We separately measured the achromatic contrast of the female wings (first row of each figure) and male wings (second row of each figure) to account for sexual dimorphism. The dotted line represents the threshold of discrimination by any visual model: visual discrimination is considered possible if the measured chromatic distance is superior to the threshold. The overlapping between the confidence intervals and the discrimination threshold shows that neither a bird visual model nor a Morpho visual model could discriminate between the wing brightness of Morphos from allopatric and sympatric populations.

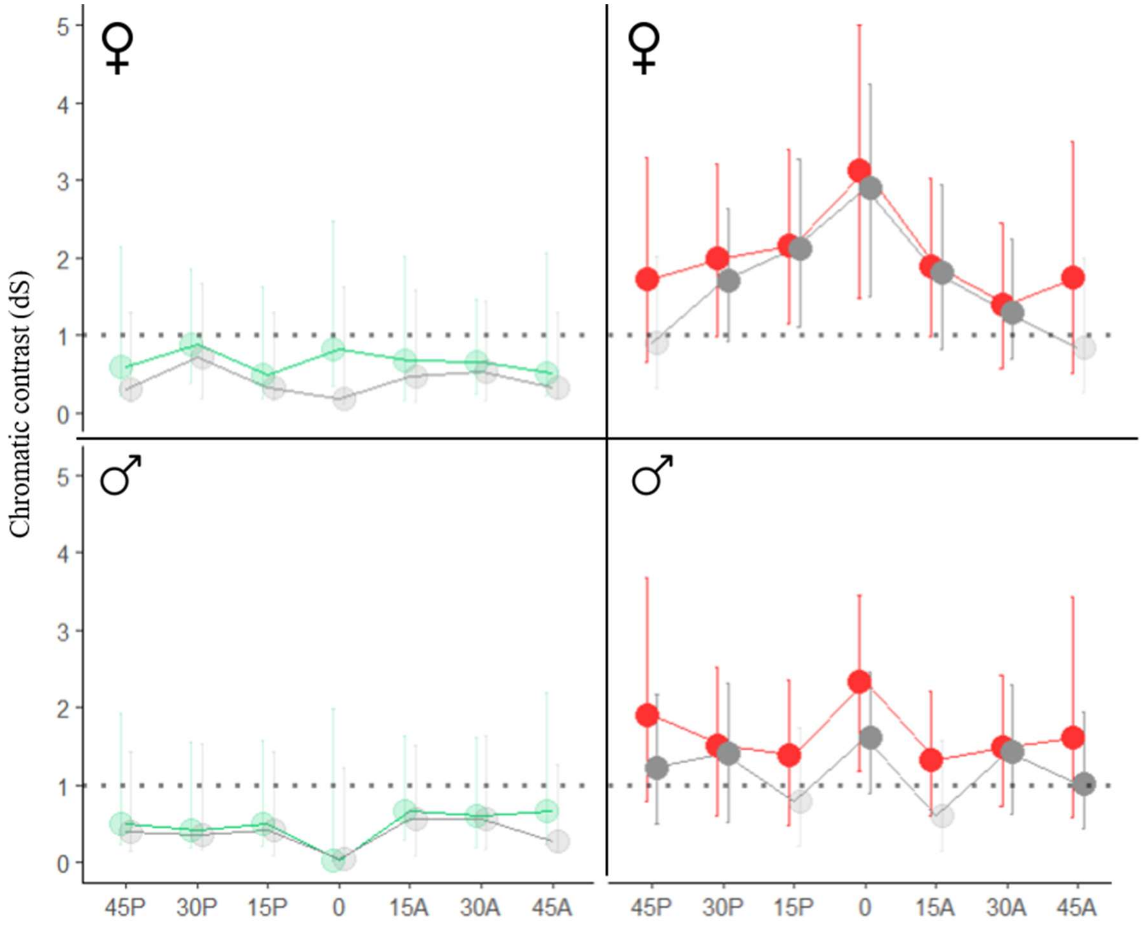


**S7 Fig** Chromatic distances (i.e. the visual discrimination rate by a visual model) of the wing reflectance measured with the "Specular" set-up on the anteroposterior plane. We used visual modeling to calculate the chromatic contrast found between the blue coloration of the two allopatric population of *M.helenor* sampled in Ecuador, as seen by a *Morpho* visual system (in red) and between the blue coloration of two species of Morphos from French Guiana, as seen by a *Morpho* visual system (in green). We also added the chromatic contrast of the *Morpho* wings as seen by the visual system of an avian predator (in grey). We separately measured the chromatic contrast of the female wings (first row) and male wings (second row) to account for sexual dimorphism. The dotted line represents the threshold of discrimination by any visual model: visual discrimination is considered possible if the measured chromatic distance is superior to the threshold. Overall, the results are similar to the chromatic distances measured on the reflectance spectra extracted from the proximo-distal plane of the *Morpho* wings (Fig. 3.C in the main text): the minute hue differences observed between species on the anteroposterior plane of the wings cannot be discriminated in a *Morpho* visual model, whereas the intraspecific divergent hues measured on the wings of allopatric *M. h. bristowi* and *M. h. theodorus* can be discriminated by a *Morpho* visual model.

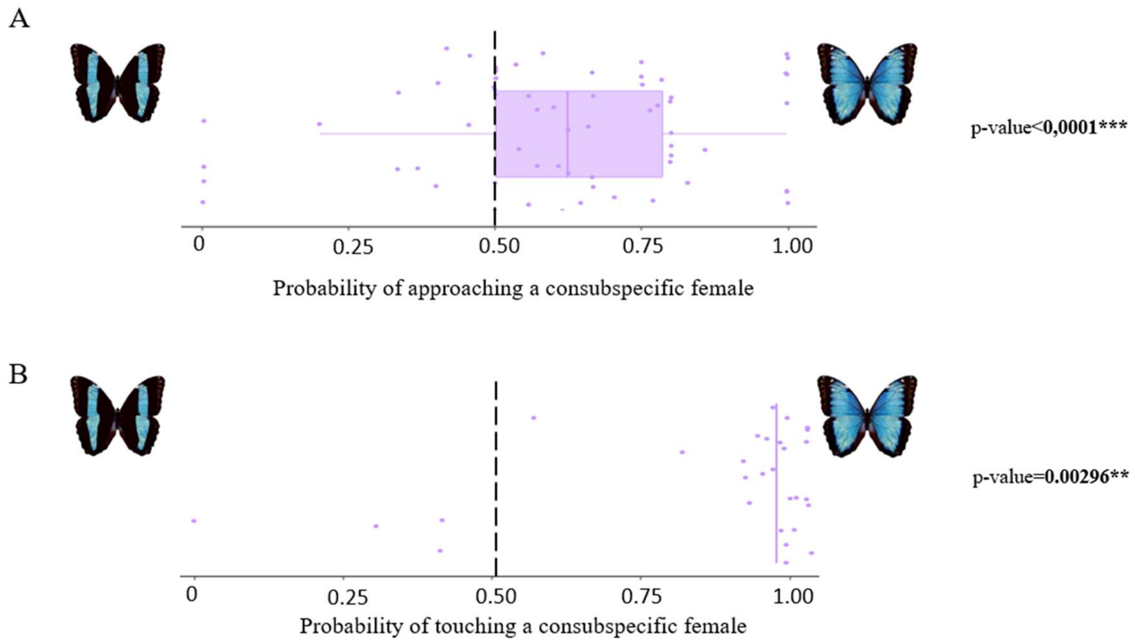


**S8 Fig** Probabilities of (A) approaching and (B) touching a con-subspecific female for *M. h. bristowi* males (purple). The *M. h. bristowi* males had the choice between a wild-type *M. h. bristowi* female model (on the right) and a modified *M. h. bristowi* female model with a narrowed blue band pattern like *M. h. theodorus* (on the left). The dotted line indicates the probability of approaching/touching a model expected if no preference was found. The p-values of Wilcoxon tests testing the significant departure from the 0.5 probability are shown next to the corresponding graphs.

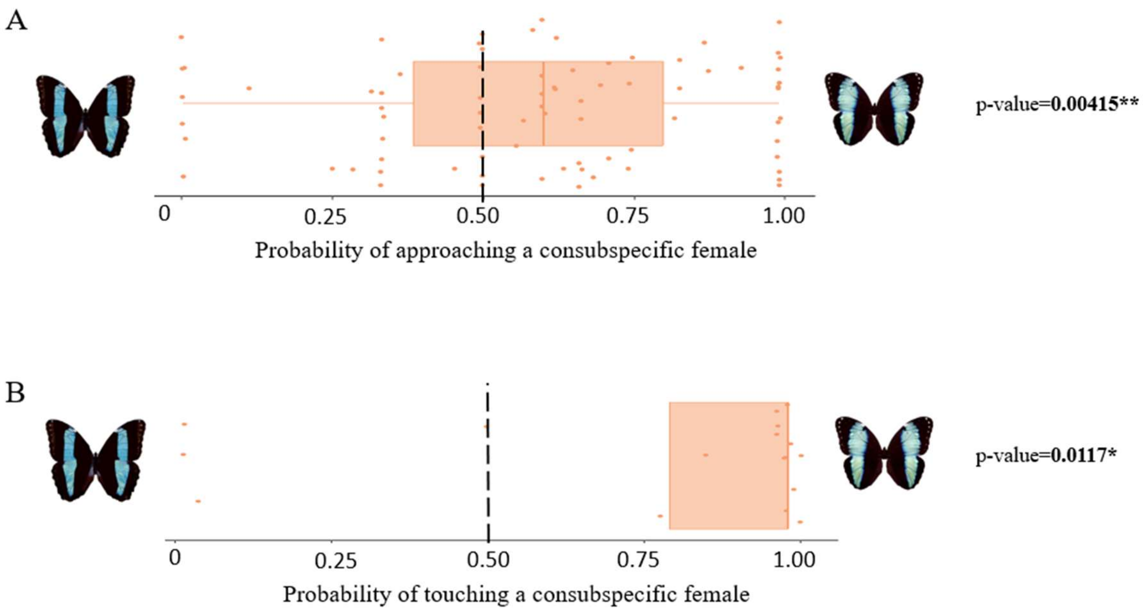


**S9 Fig** Probabilities of (A) approaching and (B) touching a con-subspecific female for *M. h. theodorus* males (orange). The *M. h. theodorus* males had the choice between a WT *M. h. theodorus* female model and a modified *M. h. bristowi* female model with a narrow blue band pattern like *M. h. theodorus*. The dotted line indicates the probability of approaching/touching a model expected if visual preferences are lacking. The p-values of Wilcoxon tests testing the significant departure from the 0.5 probability are shown next to the corresponding graphs.

**A**

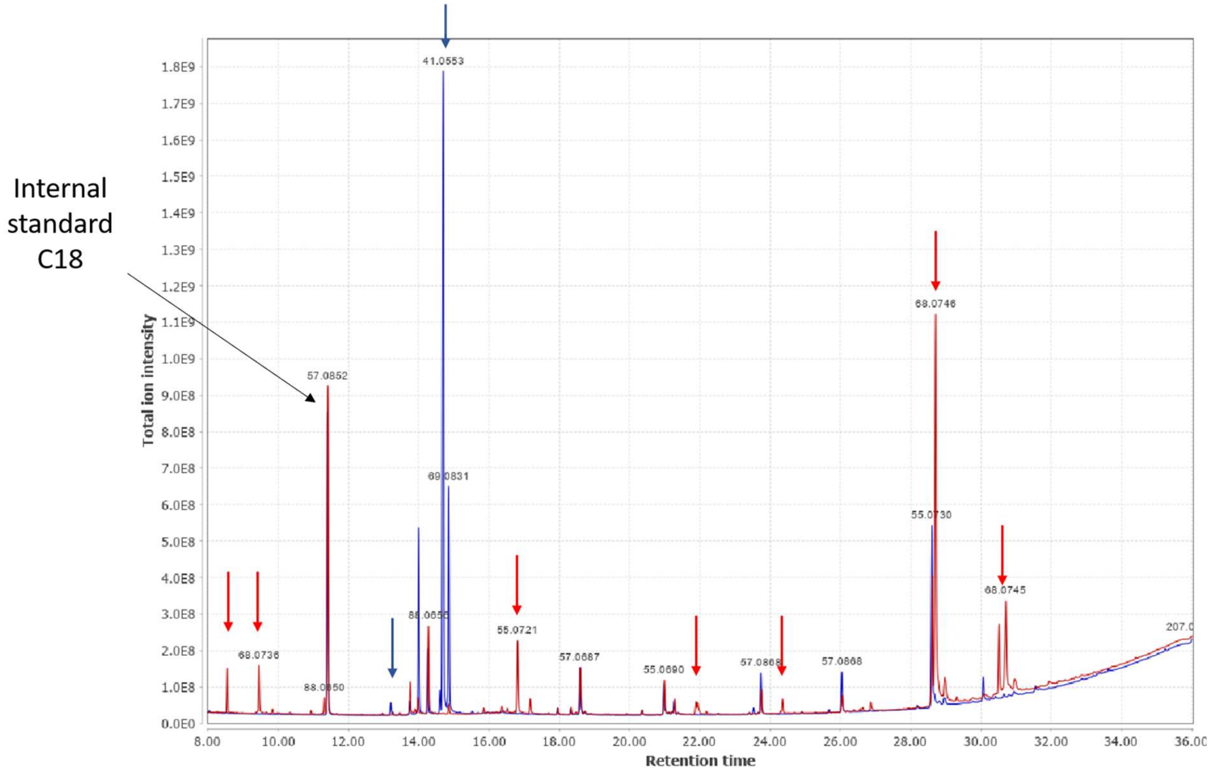

Internal standard C18
Total ion intensity
Retention time
41.0553
57.0852
68.0746
55.0730
68.0745
55.0721
57.0687
55.0690
57.0868
68.0736


**B**

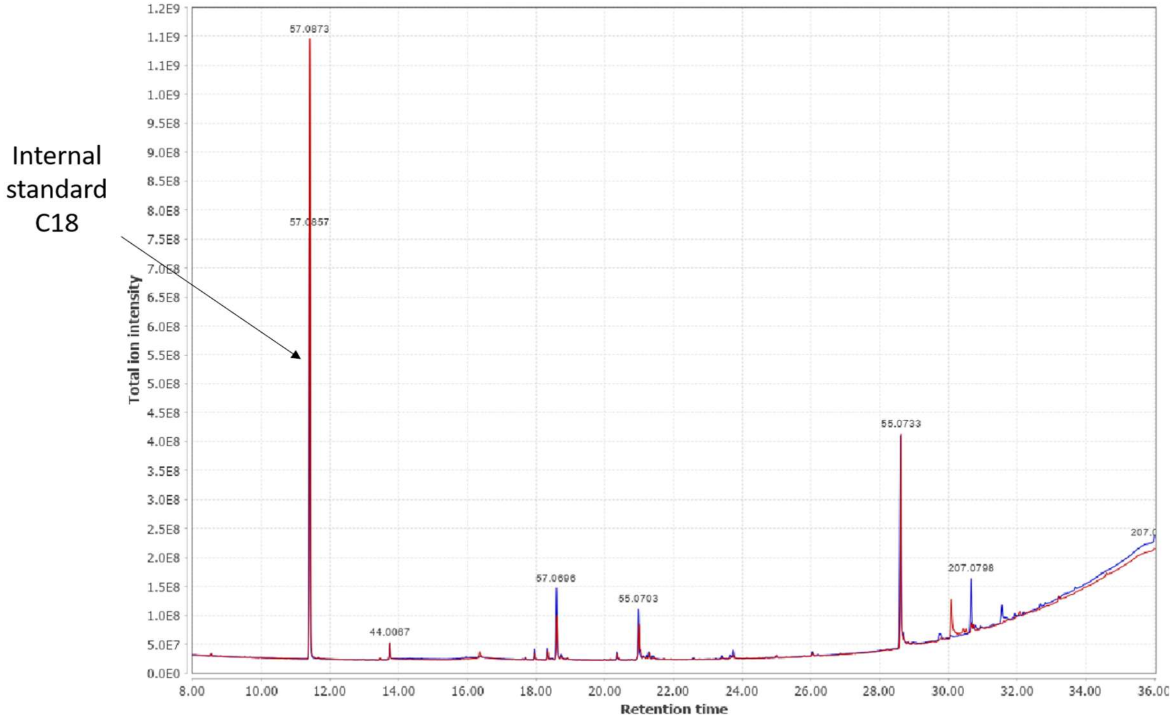

Internal standard C18
Total ion intensity
Retention time
57.0873
57.0857
55.0733
207.0798
57.0696
55.0703
44.0067

**S10 Fig** Example of chromatogram obtained when comparing the chemical compounds from C16 to C30 found on male genitalia (A) and on female genitalia (B). The red chromatogram shows the chemical compounds of *M. h. helenor* individuals and the blue one the chemical compounds of the *M. a. achilles* individuals. Arrows point at specific compounds only found in *M. h. helenor* males (red arrows) or *M. a. achilles* males (blue arrows).

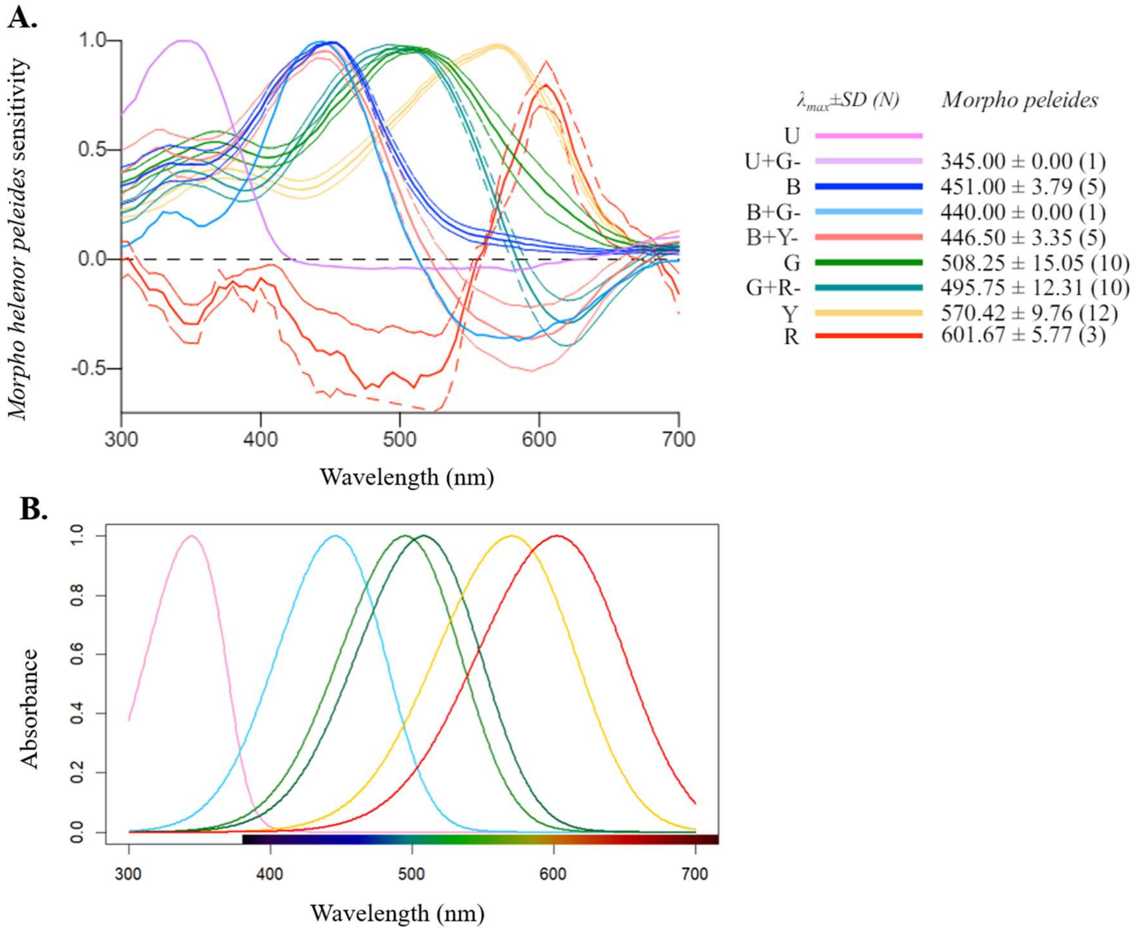


**S11 Fig** (A) *Morpho helenor* visual sensitivities from **(**2) and (B) the absorption of the cones used in our visual *Morpho* model to study the discrimination of the blue coloration in these butterflies.

**Table S1.** Permutation-based ANOVAs performed in order to test whether sex, taxa or the angle of illumination have an effect on the estimated hue and brightness measured on the proximo-distal plane of their wings. We observe that the effect of sex is always significant on the variations of brightness hinting at sexual dimorphism. Hue is also significantly different between the allopatric sub-species of *M. helenor*, *M. h. theodorus* and *M. h. bristowi*, whereas it was very similar for the two sympatric species *M. h. helenor* and *M. a. achilles*, suggesting convergence in wing coloration between the two sympatric species.

| | Hue Specular French Guiana | Hue Specular Ecuador | Brightness Specular French Guiana | Brightness Specular Ecuador | Brightness « 30° tilt » French Guiana | Brightness « 30° tilt » Ecuador |
|---|---|---|---|---|---|---|
| Sex | 0.062 | 0.001 *** | 0.002 ** | 0.001 *** | 0.001 *** | 0.001 *** |
| Subspecies | 0.188 | 0.001 *** | 0.001 *** | 0.066 | 0.506 | 0.007 ** |
| Angle | 0.001 *** | 0.001 *** | 0.001 *** | 0.001 *** | 0.001 *** | 0.001 *** |
| Sex:Subspecies | 0.030 * | 0.002 ** | 0.001 *** | 0.001 *** | 0.020 * | 0.114 |
| Subspecies:Angle | 0.485 | 0.638 | 0.188 | 0.362 | 0.006 ** | 0.050 * |

**Table S2.** Annotation of the chemical compounds significantly associated to *M. helenor* or *M. achilles*. We used an Indicator Value analysis to find the chemical compounds allowing to discriminate the males and females of *M. helenor* and *M. achilles* separately. Because we used a protocol allowing for the detection of small peaks during the MZmine analysis to not exclude "pheromone-like" molecules, our analysis is sensitive to small spectral variations. This could explain why some compounds were associated to different (but similar) MZmine detected occurrences. X refers to an undetermined double bond position or configuration (*Z* or *E*). The "Ions" column describes the spectrum with the major ion, followed by other ions in intensity order, and the underlined visible ion corresponding to the molecular ion.

| Sex | Dataset | Annotation | LRI | Ions | Species | P-value |
|---|---|---|---|---|---|---|
| Males | C8 to C16 | (*E*)-*β*-Ocimene | 1048 | 93 ; 79 ; 41 ; 136 | *M. helenor* | 0,0491 |
| | | Phenyl ethyl alcohol | 1112 | 91; 122; 65 | *M. achilles* | 0,0004 |
| | | Unknown compound 1 | 1123 | 80; 52; 73; 98; 124 | *M. helenor* | 0,0311 |
| | | Ethyl octanoate | 1195 | 88; 57; 101; 43; 73; 127; 172 | *M. helenor* | 0,0001 |
| | | Tetradec-1-ene | 1390 | 43; 55; 69; 83; 97; 196 | *M. achilles* | 0,0282 |
| | | Ethyl decanoate | 1394 | 88; 101; 43; 73; 155; 200 | *M. helenor* | 0,0001 |
| | | Dodec-x-enol | 1454 | 55; 68; 41; 82; 96; 184 | *M. helenor* | 0,0002 |

| | | | | | | |
|---|---|---|---|---|---|---|
| | | Ethyl dodecanoate | 1593 | 88; 101; 43; 73; 155; 228 | *M. helenor* | 0,0001 |
| | | (x)-Ethyl Tetradec-x-enoate | 1764 | 88; 55; 41; 96; 166; 254 | *M. helenor* | 0,0001 |
| | | (x)-Tetradec-x-enyl acetate | 1784 | 43; 68; 54; 82; 96; 194 | *M. helenor* | 0,0001 |
| | C16 to C30 | Ethyl hexadecanoate | 1993 | 88; 101; 43; 73; 155; 284 | *M. helenor* | 0,0001 |
| | | Unknown compound 2 | 2046 | 44; 207; 49; 83; 55; 69 | *M. helenor* | 0,0001 |
| | | Geranyl decanoate | 2148 | 69; 93; 41; 121; 136; 308 | *M. helenor* | 0,0001 |
| | | (*Z*)-Ethyl Octadec-9-enoate | 2168 | 55; 41; 69; 88; 83; 310 | *M. helenor* | 0,0001 |
| | | Ethyl octadecanoate | 2193 | 88; 101; 43; 55; 157; 312 | *M. helenor* | 0,0001 |
| | | Unknown compound 3 | 3109 | 44; 207; 57; 71; 85; 281 | *M. achilles* | 0,0096 |
| | | (x)-Tetradec-x-enyl hexadecanoate | 3148 | 68; 82; 96; 194; 43; 450 | *M. helenor* | 0,0001 |
| | | Unkown compound 4 | 3342 | 207; 43; 55; 73; 81; 95 | *M. achilles* | 0,0159 |

| | | | | | | |
|---|---|---|---|---|---|---|
| | | (x)-Tetradec-x-enyl octadecanoate | 3350 | 68; 82; 96; 194; 43; 57 | *M. helenor* | 0,0001 |
| Females | C8 to C16 | Dec-1-ene | 990 | 56; 41; 70; 83; 97; 140 | *M. helenor* | 0,028 |
| | | Dodec-1-ene | 1190 | 43; 55; 69; 83; 97; 168 | *M. helenor* | 0,0302 |

**Table S3.** Repeatability (R) of the measurements of iridescence performed on the wings of *M. h. bristowi*, *M. h. theodorus*, *M. h. helenor* and *M. a. achilles*. The repeatability was measured for the values of Brightness, Hue and Chroma at every angle of illumination.

| R | SE | Emp.2.5% | Emp.97.5% | P-value | Variable | Angle_ID |
|---|---|---|---|---|---|---|
| 0,93195387 | 0,01307307 | 0,90333517 | 0,95236442 | 4,38E-77 | Chroma | 0 |
| 0,8815925 | 0,02194644 | 0,82807312 | 0,91642739 | 2,02E-58 | Brightness | 0 |
| 0,93731446 | 0,01264365 | 0,90649166 | 0,95552368 | 7,07E-80 | Hue | 0 |
| 0,81478231 | 0,03228571 | 0,73922202 | 0,86625459 | 1,06E-43 | Chroma | 15A |
| 0,67046172 | 0,05265538 | 0,54844062 | 0,75544574 | 1,20E-25 | Brightness | 15A |
| 0,91021841 | 0,01686965 | 0,87067891 | 0,93479852 | 1,06E-67 | Hue | 15A |
| 0,77559976 | 0,03806848 | 0,68813618 | 0,83531032 | 1,52E-37 | Chroma | 15E |
| 0,6708999 | 0,05014185 | 0,56249193 | 0,74961219 | 1,10E-25 | Brightness | 15E |
| 0,87101943 | 0,02490225 | 0,81147963 | 0,90678958 | 1,41E-55 | Hue | 15E |
| 0,83143394 | 0,02937952 | 0,76803356 | 0,87725532 | 9,15E-47 | Chroma | 15I |
| 0,73512952 | 0,04070802 | 0,63907404 | 0,8005203 | 2,51E-32 | Brightness | 15I |
| 0,91839526 | 0,01578924 | 0,88039618 | 0,94433766 | 6,33E-71 | Hue | 15I |

| | | | | | | |
|---|---|---|---|---|---|---|
| 0,79982088 | 0,0350237 | 0,7162314 | 0,85456168 | 3,39E-41 | Chroma | 15P |
| 0,7198958 | 0,04867878 | 0,61651339 | 0,79526635 | 1,36E-30 | Brightness | 15P |
| 0,88930422 | 0,02020534 | 0,8443417 | 0,92331442 | 1,14E-60 | Hue | 15P |
| 0,79421599 | 0,03598134 | 0,7191763 | 0,8551282 | 2,62E-40 | Chroma | 30A |
| 0,67732938 | 0,04919037 | 0,56751756 | 0,76016964 | 2,80E-26 | Brightness | 30A |
| 0,9289188 | 0,01305597 | 0,89974257 | 0,95162324 | 1,33E-75 | Hue | 30A |
| 0,70444585 | 0,04668047 | 0,60669801 | 0,78777167 | 6,07E-29 | Chroma | 30E |
| 0,71178183 | 0,045488 | 0,61393897 | 0,79154588 | 1,03E-29 | Brightness | 30E |
| 0,87874635 | 0,02331122 | 0,82296662 | 0,91409894 | 1,25E-57 | Hue | 30E |
| 0,6550905 | 0,05264239 | 0,53816018 | 0,74507705 | 2,76E-24 | Chroma | 30I |
| 0,74554083 | 0,04261923 | 0,64566456 | 0,81391148 | 1,41E-33 | Brightness | 30I |
| 0,87582059 | 0,02347792 | 0,82228053 | 0,91484364 | 7,75E-57 | Hue | 30I |
| 0,9041886 | 0,01878598 | 0,85773967 | 0,93360667 | 1,64E-65 | Chroma | 30P |
| 0,81205069 | 0,03280043 | 0,74302844 | 0,86991471 | 3,15E-43 | Brightness | 30P |
| 0,97112453 | 0,00585086 | 0,95746447 | 0,9795759 | 1,88E-106 | Hue | 30P |
| 0,85400673 | 0,0260558 | 0,79757803 | 0,89737906 | 1,77E-51 | Chroma | 45A |
| 0,77039882 | 0,0374088 | 0,69002671 | 0,83570177 | 8,13E-37 | Brightness | 45A |

| | | | | | | |
|---|---|---|---|---|---|---|
| 0,90203576 | 0,01936148 | 0,85524126 | 0,93076324 | 9,14E-65 | Hue | 45A |
| 0,75913134 | 0,03786569 | 0,67916231 | 0,82351654 | 2,66E-35 | Chroma | 45E |
| 0,81757629 | 0,02896174 | 0,75431007 | 0,87015713 | 3,40E-44 | Brightness | 45E |
| 0,82071206 | 0,03200376 | 0,74873736 | 0,86827257 | 9,31E-45 | Hue | 45E |
| 0,74259191 | 0,04170908 | 0,64865071 | 0,80893911 | 3,23E-33 | Chroma | 45I |
| 0,71182855 | 0,04478549 | 0,62281411 | 0,7850276 | 1,02E-29 | Brightness | 45I |
| 0,84104241 | 0,02886878 | 0,77925018 | 0,88941637 | 1,10E-48 | Hue | 45I |
| 0,83345745 | 0,03061064 | 0,76221461 | 0,88316779 | 3,69E-47 | Chroma | 45P |
| 0,72576245 | 0,0435171 | 0,62298039 | 0,79267106 | 3,01E-31 | Brightness | 45P |
| 0,79537205 | 0,03348855 | 0,72196461 | 0,85099538 | 1,73E-40 | Hue | 45P |
| 0,76096223 | 0,03875962 | 0,67466542 | 0,82646812 | 1,53E-35 | Chroma | 30A+ |
| 0,57007786 | 0,05936361 | 0,44073031 | 0,68032181 | 6,32E-18 | Brightness | 30A+ |
| 0,82901303 | 0,03012831 | 0,75897509 | 0,87566813 | 2,67E-46 | Hue | 30A+ |
| 0,89830354 | 0,01859355 | 0,85566402 | 0,9260342 | 1,65E-63 | Chroma | 30E+ |
| 0,60590647 | 0,05531483 | 0,48615964 | 0,6922734 | 2,16E-20 | Brightness | 30E+ |
| 0,86040869 | 0,02652343 | 0,79782212 | 0,90284859 | 5,85E-53 | Hue | 30E+ |
| 0,83151877 | 0,03001095 | 0,76702206 | 0,87982723 | 8,82E-47 | Chroma | 30I+ |

| | | | | | | |
|---|---|---|---|---|---|---|
| 0,80264773 | 0,03329181 | 0,72607819 | 0,85682716 | 1,18E-41 | Brightness | 30I+ |
| 0,75740508 | 0,03975858 | 0,67291885 | 0,81765625 | 4,46E-35 | Hue | 30I+ |
| 0,83149921 | 0,03123307 | 0,76071749 | 0,88420299 | 8,89E-47 | Chroma | 30P+ |
| 0,7315642 | 0,0418415 | 0,64205063 | 0,80411359 | 6,54E-32 | Brightness | 30P+ |
| 0,54180507 | 0,06347357 | 0,39773273 | 0,64264721 | 3,63E-16 | Hue | 30P+ |
| 0,82564933 | 0,03141051 | 0,75785206 | 0,87820262 | 1,15E-45 | Chroma | 30A- |
| 0,70304331 | 0,04565709 | 0,609797 | 0,78119994 | 8,47E-29 | Brightness | 30A- |
| 0,70430795 | 0,04743961 | 0,59725879 | 0,77909707 | 6,27E-29 | Hue | 30A- |
| 0,76953439 | 0,04063247 | 0,68300252 | 0,83811216 | 1,07E-36 | Chroma | 30E- |
| 0,84800758 | 0,02819233 | 0,78156911 | 0,88972866 | 3,73E-50 | Brightness | 30E- |
| 0,84527977 | 0,0268152 | 0,7854896 | 0,88880349 | 1,43E-49 | Hue | 30E- |
| 0,9142409 | 0,01566538 | 0,87865615 | 0,93819666 | 3,01E-69 | Chroma | 30I- |
| 0,83297497 | 0,02802545 | 0,76998544 | 0,88057836 | 4,59E-47 | Brightness | 30I- |
| 0,88792846 | 0,02111732 | 0,84024728 | 0,92021421 | 2,95E-60 | Hue | 30I- |
| 0,7596208 | 0,04114026 | 0,6698537 | 0,826702 | 2,29E-35 | Chroma | 30P- |
| 0,7115446 | 0,04568159 | 0,61202912 | 0,78354985 | 1,09E-29 | Brightness | 30P- |
| 0,9412519 | 0,01089593 | 0,91657877 | 0,95702933 | 4,35E-82 | Hue | 30P- |

**S1 Dataset (separate file):** All raw measurements of wing reflectance and R scripts required to generate the results are available under the DOI 10.5281/zenodo.14389631 https://doi.org/10.5281/zenodo.14389631
